# CRISPRs are associated with bacterial expansion in the human gut during the first year of life

**DOI:** 10.1101/2021.09.03.458793

**Authors:** Jiqiu Wu, Timothy G. Barraclough

## Abstract

The dynamics of the gut microbiome in infancy have a profound impact on health in adulthood, but the ecological mechanism underlying the dynamics between bacteria and bacteriophages remains poorly understood. CRISPR is a bacterial adaptive immune system to resist bacteriophages; however, the role they play in the dynamics in infants’ gut microbiota is unknown. In this work, using large-scale metagenomic sequencing data from 82 Sweden infants’ gut microbiomes, 1882 candidate CRISPRs were identified and their dynamics were analyzed. The results showed CRISPRs were distributed in dominant bacteria and could target distinct bacteriophages at different time points with largely alternated spacers. In the putative identical CRISPRs, we found the CRISPRs could acquire new spacers and loss old spacers during the first year. In addition, it is the first time to report that gender was a major factor to determine the bacterial richness and the number of CRISPRs and host range of bacteriophages was narrow *in silico*. Therefore, we concluded that CRISPRs were associated with bacterial expansion. This work improves the understanding of the ecological mechanism behind dynamics in the early life of the human gut and substantially expands the repertoire of predicted CRISPRs providing a resource to study the function of bacterial unknown genes and to enhance the performance of these beneficial bacteria by CRISPR gene-editing technology.

## 1 Introduction

The rise of metagenomic sequencing has uncovered the high diversity of bacteria, archaea, fungi, and viruses living in human gut microbiomes (1). These microbes have been associated with a great many diseases, such as cardiovascular disease (2), depressive disorder (3), and obesity (4). In particular, the microbiota of infants is a key factor for the development of the microbiome in adulthood (5) and has a significant effect on basic neurogenerative processes (6) and immune system development (7). In previous studies, cessation of breast-feeding, mode of delivery (8), geographical location and household exposures (5) were identified as major factors in determining the dynamics of infants’ gut microbiota. Yet, the underlying ecological mechanism behind such changes had not been studied.

One possible source of variation in the composition of microbiomes is the interaction between bacteria and bacteriophage in the gut community. Lim et al (9) showed that the expansion of the bacterial community during the first year of life is accompanied by a contraction of the bacteriophage community. However, the ecological mechanisms and nature of interactions between bacteria and viruses has not been widely investigated. The emergence and dynamics of bacterial resistance mechanisms to bacteriophages might be critical to this process.

CRISPRs, clustered regularly interspaced short palindromic repeats (10) are an important adaptive immune system found in most archaea and many bacteria for resistance to phages (11). Although the organization of CRISPR locus is very diverse (12), generally, it consists of two main components: the CRISPR array whose function is genetic memory, and the Cas (CRISPR-associated) proteins, which is the catalytic core of the system. The CRISPR array is composed of repetitive sequences, repeats, that are separated by variable sequences, spacers. The spacer sequences are derived from invading mobile genetic elements, such as bacteriophages (13). The crRNAs encoded by the spacers guide the complexes of Cas proteins to the complementary bacteriophage target sequences that match the spacers. Cleavage of the sequences is then catalyzed by various Cas enzymes (14).

There have been some studies on CRISPRs of single bacterial species and genera, such as *Streptococcus thermophilus* (15), *Streptococcus* (16), and *Bifidobacterium longum* (17), demonstrating that CRISPRs enhance phage resistance by a process of spacer expansion and loss. Nevertheless, few studies focused the CRISPRs at a community level.

Here, we used metagenomic sequencing data to survey the whole bacterial community of 82 Swedish infants from birth to 12 months old (8), a key period during which microbiomes influencing health in later life. We analyzed how CRISPRs changed in their gut microbiomes. Our results showed that the number of spacers and CRISPRs increased with the expansion of the bacterial community with infants age. At the same time, the bacteria containing CRISPRs and the bacteriophages targeted by spacers were distinct at different time points, as a result of spacer loss and expansion. Thus, these findings indicate that CRISPRs were active and associated with bacterial expansion by combating different bacteriophages using largely alternated spacers. In addition, the community size, the number of CRISPRs and spacers of boys were significantly greater than those of girls. Taken together, we concluded that CRISPRs were associated with bacterial expansion by alternating spacers that can target distinct bacteriophages at different times. Therefore, this work enhances understanding of the dynamics of gut microbiome in early life and expands the repository of useful CRISPRs.

## 2 Methods

### 2.1 Metagenomic dataset

All the metagenomic sequencing reads data used in this research were downloaded from GigaDB (http://gigadb.org/dataset/100145, downloaded on May 20^th^ - 27^th^ in 2019) generated by an infant gut microbiome study (8). The authors extracted the DNA, constructed the library, and shotgun-sequenced stool samples from 98 mothers at delivery and their infants sampled longitudinally during the first days of life and at 4 and 12 months of age (8). In this research, some subjects with incomplete metadata were filtered out, such as those missing antibiotic usage and feeding pattern (see metadata in Additional file 1 in Supplementary Information).

### 2.2 *De Novo* assembly of contigs

Adaptor contamination, low-quality reads, and host contaminating reads were removed from the raw sequencing reads by Ba ckhed and his colleagues (8). The clean reads were assembled *de novo* using MEGAHIT (18) with the option *–no-mercy*, which recovers the sequences at very low depth and strengthens the contiguity of low-depth regions for metagenomics assembly. It is important here to recover a taxonomically representative sample, as many bacteria will be present at low frequency. The community size was defined as the sum of the base-pairs in all assembled contigs based on the MEGAHIT output and it described the total number of the detected nucleotides in the community.

### 2.3 Bacterial metagenomics analysis

At the whole-metagenome level, we characterized the taxonomic profiles by MetaPhlAn2 (19) with the default parameters, which searches for unique clade-specific marker genes. Bacterial richness (α-diversity) was calculated based on the output of MetaPhlan2. To investigate changes in bacterial composition over time, principal coordinates analysis (PcoA) was performed in the *vegan* package in R on Bray–Curtis dissimilarities as a measure of β-diversity. To judge whether sequencing depth was adequate to infer composition, the *ape* package was used to conduct rarefaction curves with 500 permutations. Additionally, the difference of relative abundance at genus level and the bacterial richness between boys and girls was compared by calculating *P* values using Mann-Whitney U test in R, due to the variances were not normally distributed.

### 2.4 Detection of CRISPR arrays

CRISPR arrays were detected by CRISPRCasFinder (20) based on “repeat – spacer” like structure with the default parameters. CRISPRCasFinder can detect the shortest CRISPR structures, but the background of spurious candidates can be very high (21). CRISPRCasFinder includes a rating system based on several criteria to discriminate spurious CRISPR-like elements from CRISPR candidates. CRISPR arrays with evidence-levels 3 and 4 were considered as highly likely candidates, whereas evidence-levels 1 and 2 indicate potentially invalid CRISPR arrays. In addition, recent evidence suggests that isolated CRISPRs that lack Cas genes in its vicinity region can be non-functional (orphan), or work with a distant Cas locus in the same genome (22), which is hard to identify with metagenomic sequencing data. Therefore, we restricted analysis to CRISPRs with evidence level ⩾3 and that contain at least one Cas protein. The density of CRISPRs was defined as the number of CRISPRs divided by the community size after assembly and it described how many of all nucleotide bases in a community are CRISPRs.

### 2.5 Taxonomic assignments of bacterial genome and phage target of each CRISPR

Contigs with CRISPRs were queried against the NCBI NT database using BLASTn (23) (*e*-value cutoff: 1E−5, word size cut off: 100, and identity percentage: 70) in order to identify the bacterial species containing each CRISPR. Spacers were queried against a customized viral database downloaded from NCBI website (ftp://ftp.ncbi.nlm.nih.gov/refseq/release/viral/, downloaded on 1st January 2019) using BLASTn for short task (*e*-value cutoff: 1E-5, bit scores: 50, which roughly correlates to 2-nt differences over the 30-nt average length of the spacers (24)) in order to identify the bacteriophages targeted by the spacers.

### 2.6 Interaction analysis between bacteria and bacteriophages

To infer the interaction between known bacteria and known bacteriophages, the CRISPRs with spacers whose bacteriophage targets were known were used to detect the bacterial host. The phylogenetic trees of bacteriophages and bacteria were built by VICTOR (25) and TimeTree (26), respectively. The interaction trees were built using *ape* package in R.

### 2.7 CRISPR array analysis and spacer analysis

The number of candidate CRISPRs and spacers in the gut microbiota of each infant at different times was summarized. Analysis of variance (ANOVA) was carried out on the factors affecting the number of CRISPRs and spacers, such as gender, age, antibiotics usage, mode of delivery, and cessation of breast-feeding in R.

The bacteria that contain CRISPRs at phylum and genus level were compared between boys and girls using Mann-Whitney U test in R. Spacers shared between different time points, the bacteria that contain the shared spacers, and the bacteriophage that the shared spacers target were identified. The number of phylum and genus of bacterial genome that spacer appeared was calculated. All the plots were visualized using ggplot2 package in R.

### 2.8 Longitudinal analyses of the putative identical CRISPR array

In order to study the continuous changes of spacer in the identical CRISPR array, putative cases of identical CRISPR sequenced at multiple time points were selected based on the following criteria: They are detected in (1) the same infant but at different time points and in (2) the same bacterial genome, (3) their Cas protein structure is the same and (4) the *e*-value of BLASTn is less than 1E−5. The CRISPR arrays was reconstructed manually and three groups of CRISPRs meeting these criteria were highlighted. Single nucleotide dynamics of the spacers in putative identical CRISPR array was analyzed by comparing the similarity of the unique spacers.

## 3 Results

### 3.1 The number of CRISPRs and spacers increased with bacterial expansion

We identified 1882 candidate CRISPRs in the assembled contigs (the quality of assemblies was shown in Fig S1) of our filtered dataset with complete metadata for 82 infants across three time points (metadata in Additional file 1, information of these candidate CRISPRs in Additional file 2, sequences of these CRISPRs in Additional file 3). The number of CRISPRs and spacers increased with age (Fig 1a – 1b; ANOVA: the number of CRISPRs: F = 108.284, df = 2, P < 2e-16; the number of spacers: F = 93.058, df = 2, P < 2e-16). However, the average number of spacers within the same CRISPR over time showed no such trend (Fig 1c) and kept constant. The density of CRISPRs (defined in Methods) kept constant as well (Fig 1d). These trends were associated with expansion of the bacterial community with age (8). Analyses of our filtered dataset confirmed an increase in both α-diversity (Fig 2a, rarefaction curve of bacterial richness was shown in Fig S2) and community size (defined in Methods) (Fig 2b) with age and the number of CRISPRs and spacers were correlated strongly with bacterial richness (Fig 2c – 2d, the number of CRISPRs: R^2^ = 0.4029, p = 2.2e-16; the number of spacers: R^2^ = 0.3687, p = 2.2e-16).

**Fig 1.**
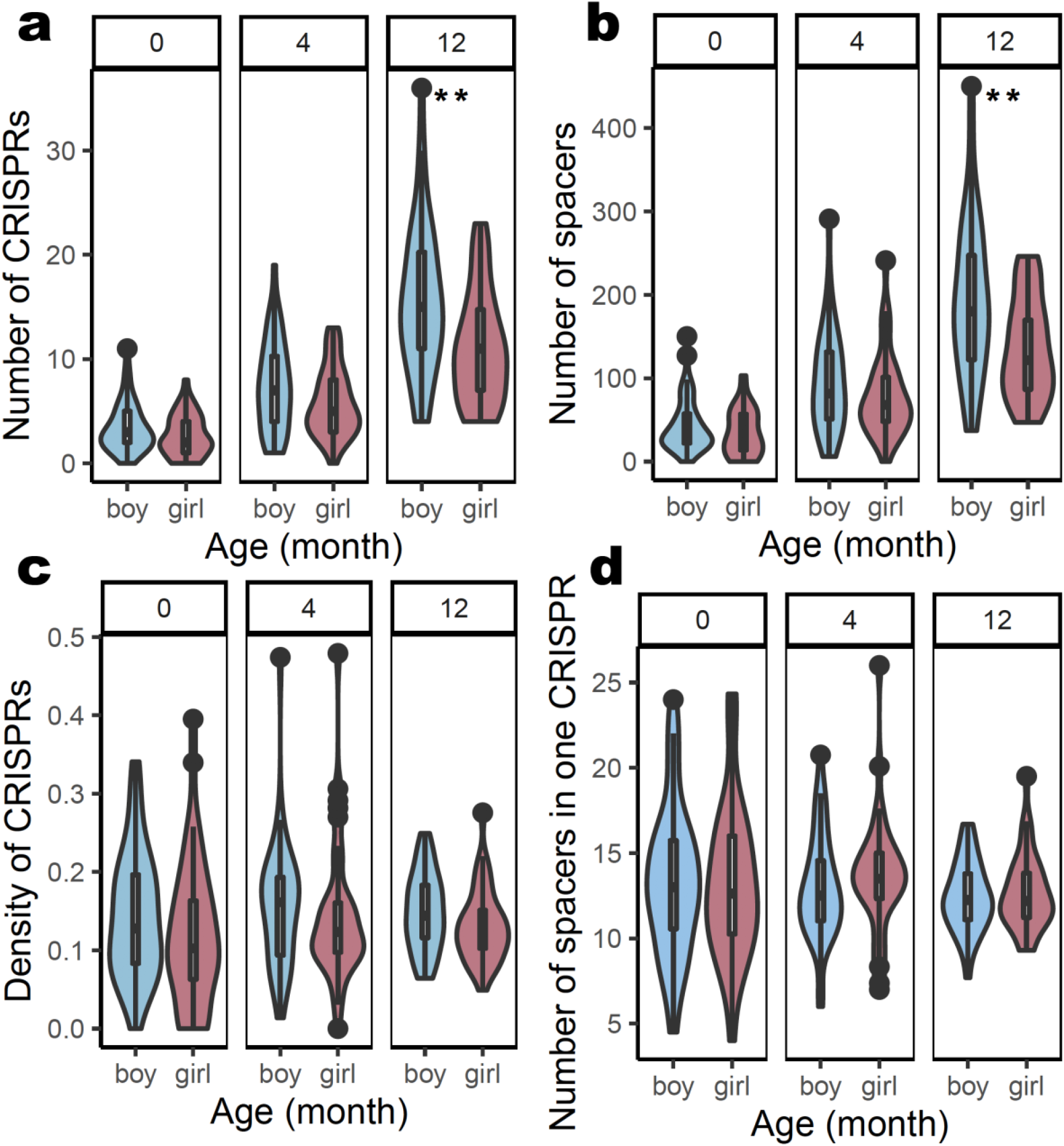
The number of CRISPRs and spacers increased with age. CRISPRs and spacers were identified in assembled contigs of gut microbiome metagenomic sequencing data. (a) and (b) showed the number of CRISPRs and spacers increased with age, respectively, and the gender difference was significant. However, the density of CRISPRs (c) and the number of spacers in one CRISPR (d) kept constant during the first year. Red color indicates girls and blue color indicates boys. *: p < 0.05, **: p <0.01, ***: p<0.001, Wilcoxon test. Sample size: boy: n = 40, girl = 42.

**Fig 2.**
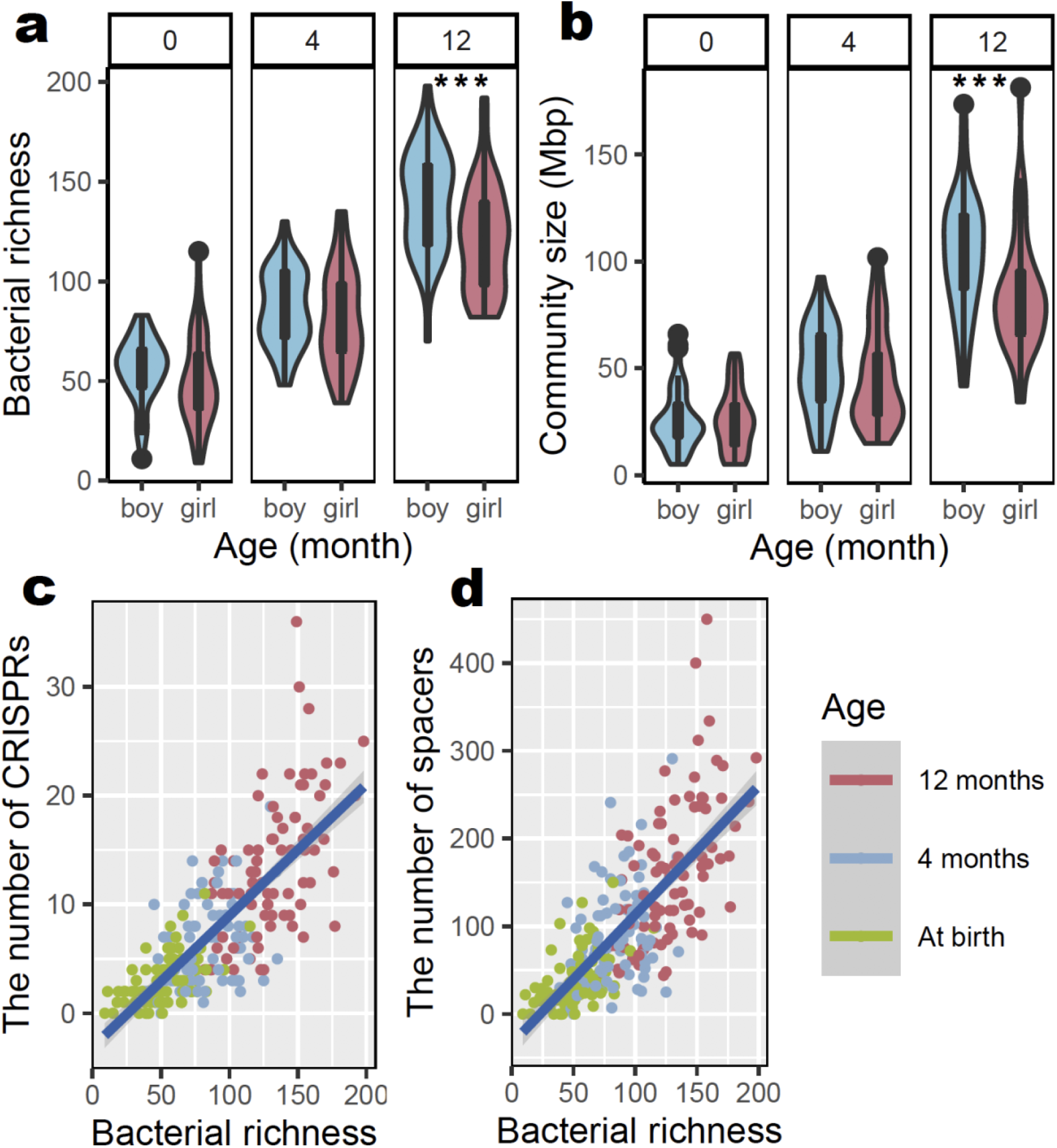
The number of CRISPRs and spacers increased with bacterial expansion. Community size and bacterial community were analyzed using MetaHIT and Metaphlan2. (a) and (b) showed bacterial expansion with the increased bacterial richness and community size during the first year. In addition, the number of CRISPRs and spacers increased with bacterial expansion. (c) and (d) showed the positive linear correlation between the number of CRISPRs and spacers with bacterial richness. Red color indicates girls and blue color indicates boys. *: p < 0.05, **: p <0.01, ***: p<0.001, Wilcoxon test. Sample size: boy: n = 40, girl = 42.

### 3.2 Spacers targeted distinct different phages at different time

We compared the sequences of spacers detected and found spacers present in CRISPR arrays changed greatly over the first year of life. Overall, 23894 spacer sequences were found and 3264 of them were found at birth, 7157 at 4 months, and 13473 at 12 months, corresponding with the increase of the number of spacers (sequences of these spacers in Additional file 4). Among them, 17269 spacers were unique, and the length of spacers mostly distributed from 30 bp to 35 bp. Interestingly, only a small fraction of spacers were shared at different time points. Among the 17269 unique spacers across the whole dataset, only 3.21% (555/17269) were shared at three time points, 4.33% (748/17269) were shared at birth and 4 month, 6.22% (1074/17269) were shared at 4 month and 12 month, and 1.12% (193/19436) were shared at birth and 12 month (Fig 3a). Consequently, within an individual as well, spacers shared at different times were scarce (Fig 3b).

**Fig 3.**
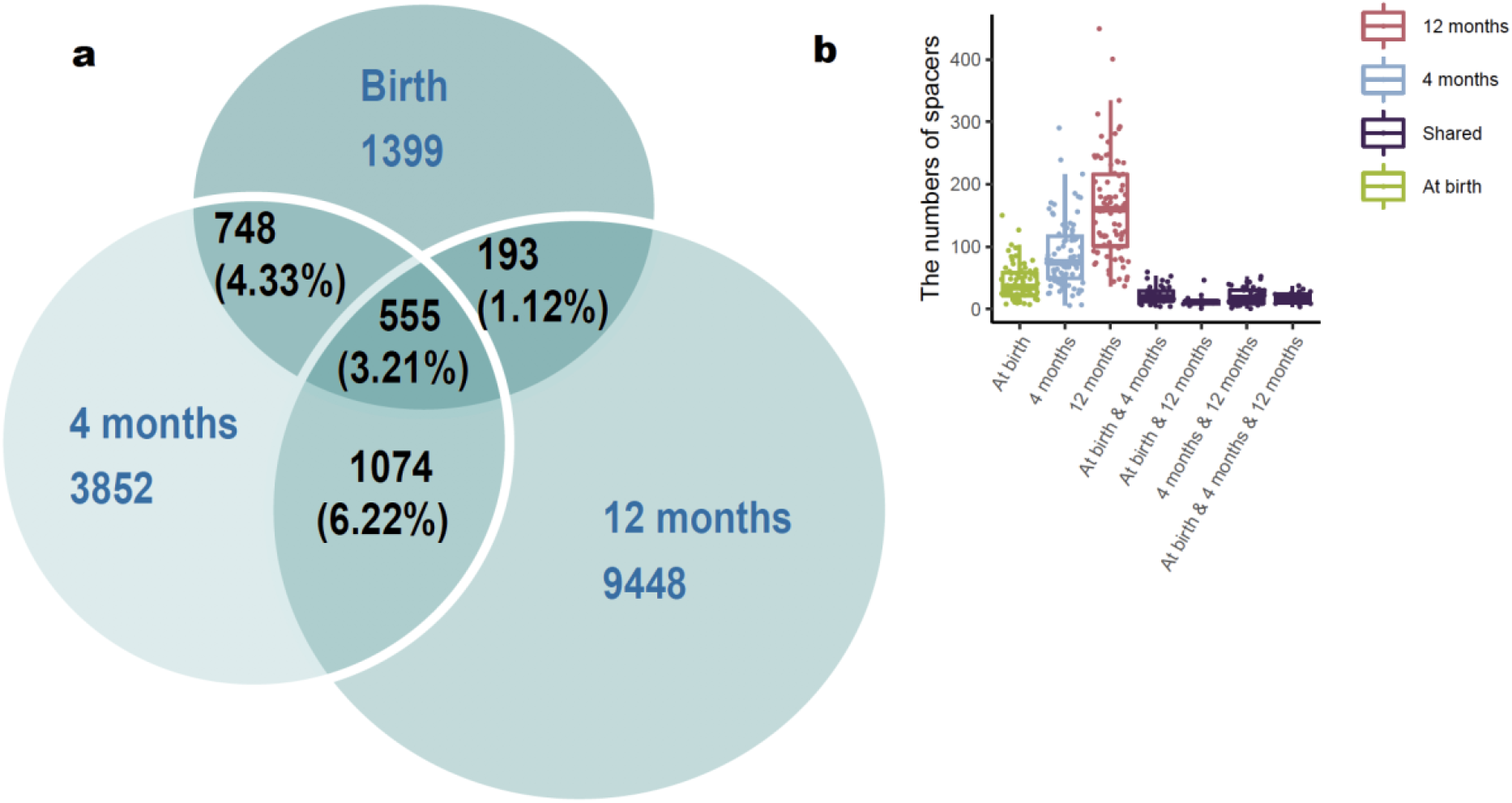
Spacers were largely alternated at different time points. (a) The Venn diagram depicts the number of shared and unique spacers at different time points among all the samples (n = 82). (b) The boxplot depicts the number of shared and unique spacers at different time points at individual level. Green color, blue color and red indicate unique spacers at birth, 4 months and 12 months, respectively. Purple color indicates the shared spacers between different time points.

Although only very few bacteriophages targeted by CRISPR could be identified by BLASTn analysis, these alternated spacers targeted very different bacteriophages during the first year. Due to limitation of the viral database, only 0.83% (161/19436) of spacers matched the virus reference genomes and most (159/19436) of the matches were bacteriophages, with 80 bacteriophages were matched during the period. Families *Myoviridae, Siphoviridae, Microviridae, Podoviridae, Herelleviridae* and *Ackermannviridae* were identified. Only *Myoviridae, Siphoviridae* and *Microviridae* were identified at all the three time points, *Podoviridae* were identified at both 4 months and 12 months, and *Herelleviridae* and *Ackmermannviridae* were only identified at 4 months and 12 months, respectively, which was consistent with rapid change of spacers between time points (Bacteriophages targeted by the spacers were shown in Fig S3). Thus, targeted phages at different times were mostly distinct.

### 3.3 CRISPRs were distributed in the dominant bacteria in the gut

After determining the bacteria host containing these CRISRPs through BLASTn, we found that CRISPRs are widely found in the dominant bacteria in the gut. Five phyla (Fig 4a) and 48 genera (Fig 4b) were included. At phylum level, Firmicutes (44.58%) and Bacteroidetes (25.43%) contained the most CRISPRs, other phyla were Actinobacteria, Proteobacteria, and Verrucomicrobia. At genus level, *Bacteroides, Bifidobacterium, Escherichia, Veillonella, and Parabacteroides* were the top 5 genera that contained the most CRISPRs.

**Fig 4.**
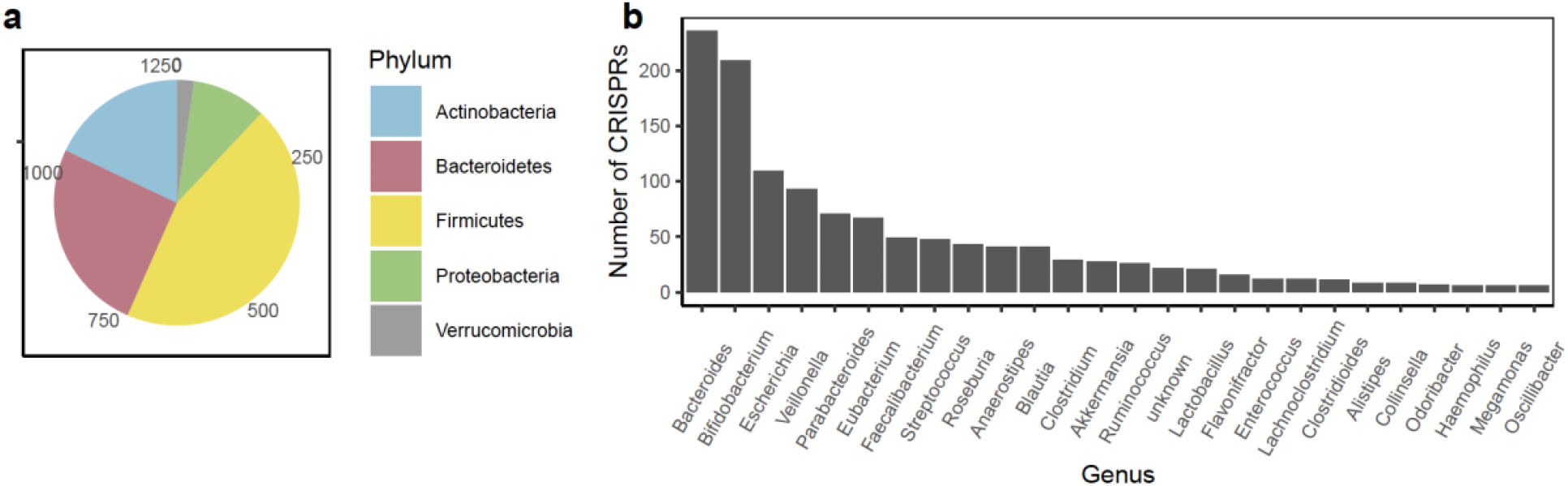
CRISPRs were distributed in dominant bacteria in gut. (a) The candidate CRISPRs were distributed in five dominant phyla in gut. (b) The candidate CRISPRs were distributed in many genera.

The interaction tree between known bacteria and bacteriophage identities showed that bacteria had CRISPR spacers for several bacteriophages, whereas a given identified bacteriophage only associates with one bacteria genus. Most of the interactions were unique and only a small part of the interactions was shared at different time points (Fig 5). In conclusion, at different time points, different dominant bacteria in the gut targeted different bacteriophages using different spacers.

**Fig 5.**
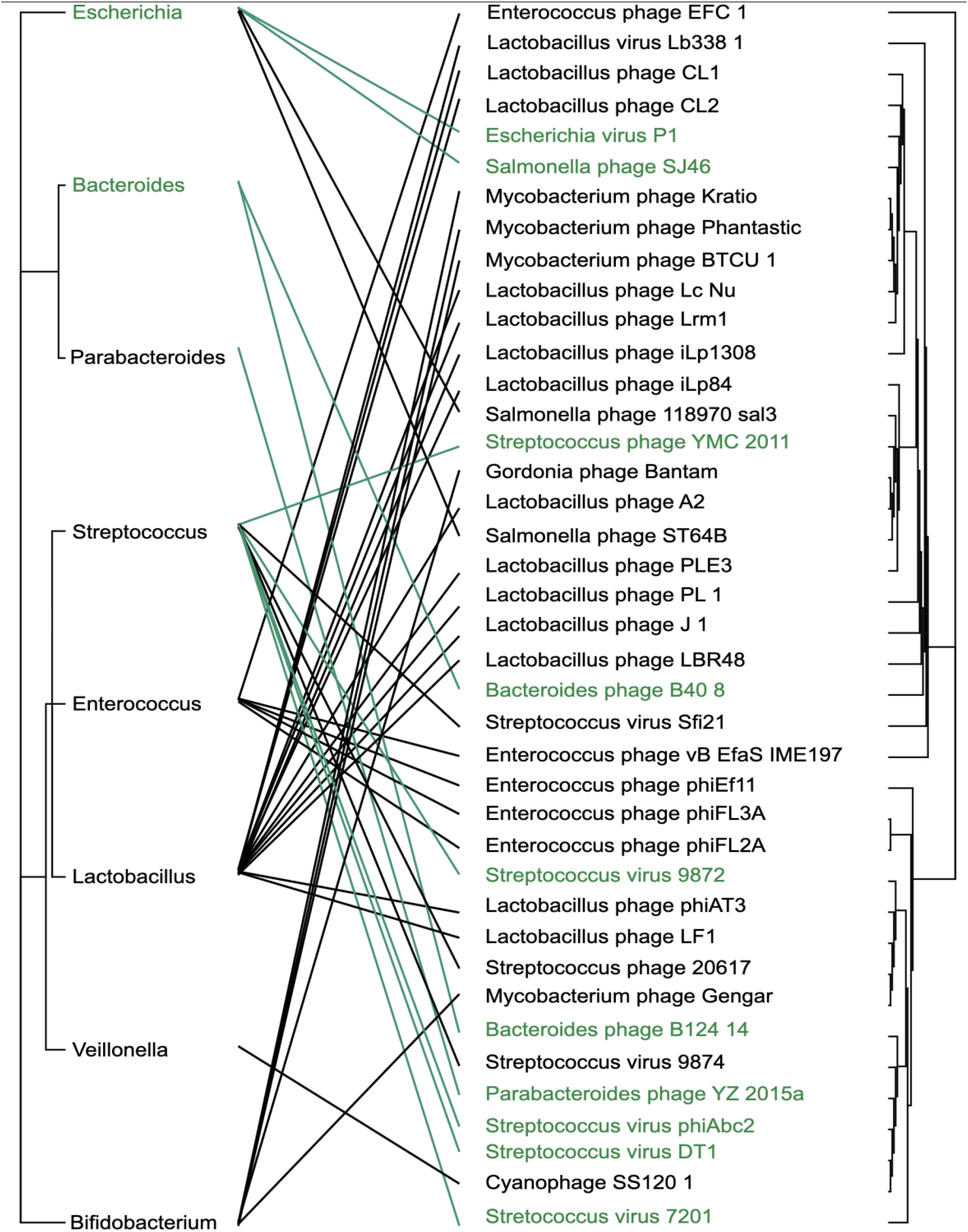
Very different bacteriophages were targeted at different time points. The interaction tree between bacteria and bacteriophages at 4 months old. The left and the right were the phylogenetic trees of bacteria and bacteriophages, respectively. The line reflected the interaction, and green indicated the unique interaction, and black indicated unique interactions at 4 months old. Interaction trees at birth and 12 months old were shown in the Fig S4.

### 3.4 CRISPRs acquired and lost spacers during the first year

To understand the dynamics in one CRISPR array, we defined putative identical CRISPR arrays (criteria see the Methods). Overall, we detected 121 putative identical CRISPR arrays (these arrays were shown in Additional file 5). Among them, spacers in 68 arrays (56.2%) changed (Fig 6a). In the changed CRISPRs, three types were observed: 31 arrays acquired new spacers, 25 arrays lost old spacers, and 12 arrays both acquired new spacers and lost old spacers (Fig 6b).

**Fig 6.**
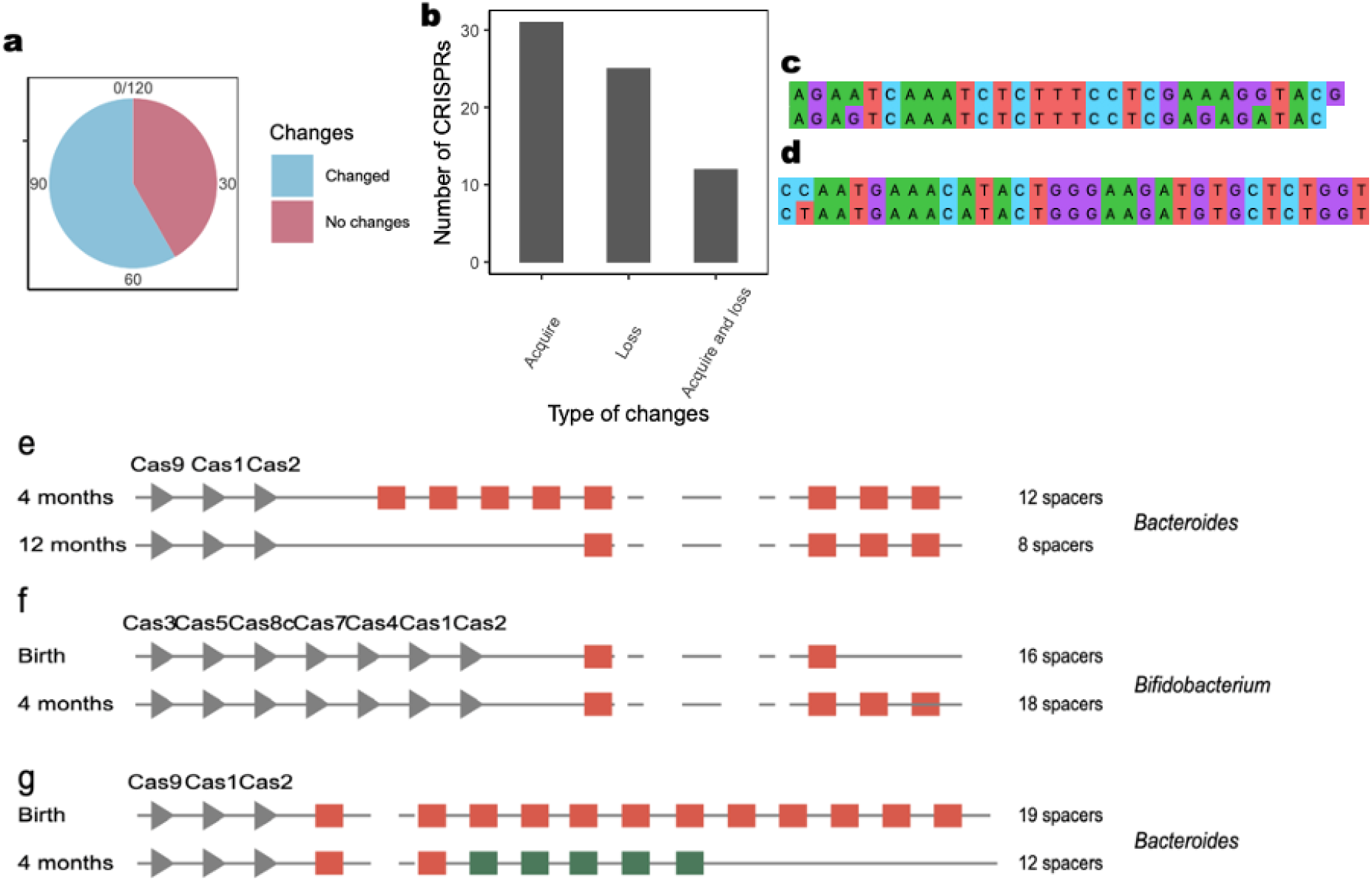
Most putative identical CRISPRs changed their arrays through acquiring and losing spacers. (a) A comparison between the changed CRISPR and the unchanged CRISPR among the putative identical CRISPRs. Blue showed the changed CRISPRs and red showed the unchanged CRISPRs. (b) showed the numbers of three types of CRISPR changes and the example of loss (e), acquire (f) and loss and acquire meanwhile (g). (c) and (d) showed that in one CRISPR, only several base differences were identified in different spacers.

Three arrays were highlighted here as examples of three types. The first array from *Bacteroides thetaiotaomicron* lost 4 spacers from 4 months to 12 months (Fig 6e). The second array from *Bifidobacterium bifidum* acquired 2 spacers from at birth to 4 months (Fig 6f). The third array from *Bacteroides vulgatus* lost 10 spacers and acquired 5 spacers from at birth to 4 months (Fig 6g). However, due to the limited reference database, we could not identify the bacteriophages that the spacers in these arrays could target.

At spacer level, longitudinal single nucleotide polymorphisms (SNP) at the same spacer position in the putative identical CRISPR was not observed. However, we found the high similarity of nucleotides of different spacers in one putative identical CRISPRs. Specifically, in the putative identical CRISPRs, we found in different spacers, there were only one or two base differences. In the first example (Fig 6c), the two spacers were in one putative identical CRISPR at 12 months from BabyID 184, the upper unique_spacer_5324 was the 5th spacer and the under unique_spacer_5328 was the 8th spacer. In the second example (Fig 6d), these two spacers were in one putative identical CRISPR at 4 months from BabyID 332, the upper unique_spacer_8429 was the 2nd spacer and the under unique_spacer_8393 was the 17th spacer.

### 3.5 Gender was the main factor that influenced the number of CRISPRs and spacers

Gender was an important factor that had effect on the number of CRISPRs and spacers by ANOVA test (Fig. 1a – 1b; ANOVA: the number of CRISPRs: F = 15.880, df = 1, p = 9.02e-05; the number of spacers: F = 13.875, df = 1, P = 0.000245). From 4 months, boys had higher number of CRISPRs and spacers than girls and when 12 months year, the difference was statistically significant (Fig. 1a – 1b, Mann-Whitney U test: the number of CRISPRs: W = 1165.5, p = 0.001261; the number of spacers: W = 1173, p = 0.001018).

To find out the reason for gender difference, we looked backwards at bacterial community data. We observed a gender difference of bacterial richness (Fig 2a) and community size (defined in Methods, Fig 2b) at 12 months (Mann-Whitney U test, bacterial richness: W = 1186.5, p = 0.0006622; community size: W = 1195, p= 0.0005034). Therefore, boys had a larger community size and higher bacterial richness than girls at 12 months.

### 3.6 Host range of bacteriophage was relatively narrow *in silico*

Previous studies have shown that the host range of bacteriophage is narrow from the experimental view. However, there are many limitations in the experiment, such as the culture of bacteria and bacteriophage, and the operation of experiments. In this study, the results showed that a small part of spacers appeared in the bacteria across species and across phylum: in total, 148 (0.76%) across genus (Fig 7a), and 2 (0.01%) spacers across phylum (Fig 7b). Therefore, the host range of bacteriophage was narrow from the metagenomic sequencing data, as well.

**Fig 7.**
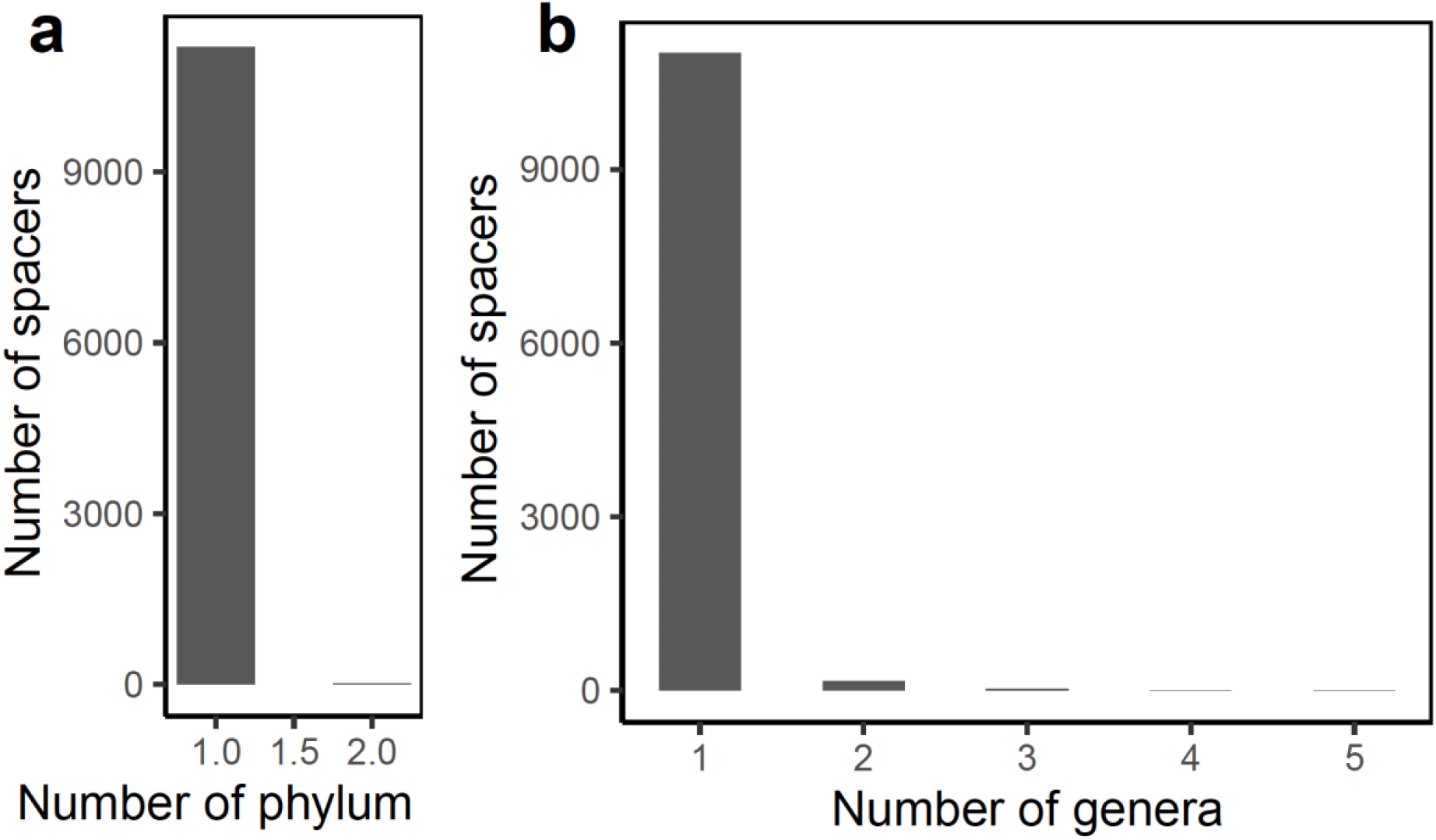
Host range of bacteriophages were narrow *in silico*. The numbers of spacers that appeared in bacteria across phylum (a) and genus (b).

## 4 Discussion

Our study analyzed the bacterial community of 82 Sweden infants’ gut microbiomes and surveyed the CRISPRs in the dynamics of bacteria and bacteriophage from birth to 12 months old. The results demonstrated the spacers in CRISPRs largely changed between time points, associated with changes in the bacteria containing CRISPRs and the bacteriophages targeted by these spacers. The 1882 candidate CRISPR systems that we identified can be used for functional mining of bacteria and to improve the bacterial ability by the gene editing technology. Moreover, it is the first time to observe the significant gender differences in the bacterial richness and the number of CRISPRs and spacers and the narrow host range of bacteriophages *in silico*.

Our results showed that CRISPRs might play a significant role during the process. Overall, the number of CRISPRs and spacers increased with bacterial expansion with infants age. More importantly, spacers, are used to target specific bacteriophages, have undergone a large-scale alternation. Spacer sharing was rare at both overall cohort level and the individual level. Thus, the bacteria containing CRISPRs and the bacteriophages targeted by spacers are distinct at different time points.

Therefore, we speculated three sources of the bacteriophages. Firstly, some bacteriophages have always resided in the gut. Secondly, some may have evolved rapidly and changed greatly from earlier time points (27). Thirdly, some may have been colonized from the environment, such as breast milk (28). On the other hand, there have been bacterial residents in the gut all the time. Meanwhile, some bacteria have been adopted from a variety of resources, such as, breast milk (29), maternal body (30), diet, house dust (31) and pets (32). These new coming bacteria brought new CRISPRs and spacers.

Moreover, existing bacteria integrate new spacers into their CRISPR arrays to provide defense against bacteriophages in response and lost the useless spacer, associated with the competitive dynamics between bacteriophages and bacteria. As a result, we observed the almost constant number of spacers in one CRISPR (Fig 1d), which is possible a balance between being able to target more bacteriophages with more spacers and the cost of maintaining these spacers. Interestingly, we observed cases with high similarity of nucleotides in different spacers in the putative identical CRISPRs, which could have resulted from acquisition of two similar spacers at the same time or sequential acquisition, perhaps by a duplication event. This diversity of similar spacers might help to cope with the diversity and rapid evolution of bacteriophage genome. In this dynamics, the molecular mechanism of how bacteria detect the environmental changes, and then regulate their CRISPR acquisition or loss of spacer is quite interesting, and more research is needed to clarify it.

Surprisingly, gender is an essential factor to affect the size and richness of bacterial community and the number of CRISPRs and spacers. More specifically, boys have increased bacterial richness and community size and consequently a higher number of CRIPSRs and spacers. Gender has been neglected in previous research on bacterial communities based on relative abundance (19), but the relative abundance of several bacteria at genus level were identified that affected by gender (33). This could result from the difference of immune system between boys and girls: males are more vulnerable to the action of the pathogen than females (34). It is possible that excess bacteria and CRISPRs may compensate for male immune deficiency and help to resist pathogens, or the greater diversity of bacteria in males could be a consequence of their weaker immune system. Further researches need to be done to elucidate the cause and mechanism of this phenomenon.

A great number of studies have documented the rising problems of antibiotic-resistant bacteria (35). Bacteriophages have been widely used in Eastern Europe for therapies for decades (36) and they have been proved to be effective in the treatment of *Salmonella* (37) and *Pseudomonas aeruginosa* and can even prevent infection (38). In this study, our results showed that the majority of these bacteriophages were able to infect bacteria only on one phylum or even one genus, which is beneficial for bacteriophages to be widely used to regulate microecology precisely without influencing other commensal and beneficial bacteria. Such target tools are urgently need in the research and application in the area of microbiome.

Gene editing technology based on CRISPR systems is also widely used to study the function of bacteria (39) and improve the performance of probiotics. Some studies have pointed out that endogenous CRISPR is more efficient and convenient than exogenous CRISPR (40). For example, a recent work emphasized endogenous CRISPR–Cas systems can be repurposed to offer new features to enhance the host colonization in lactic acid bacteria in gut, which is advantageous for human health (17). The 1882 endogenous candidate CRISPR systems we identified in this study are of great significance for subsequent gene editing engineering to study the bacterial function and enhance the performance of prebiotics.

The main limitation in our research is that assembled contigs were used to detect the CRISPR systems. In this approach, part of the reads will be omitted, leading to a loss of CRISPR quantity, as the repeat-spacer structure of CRISPR will increase the difficulty of assembly. For further research, deep sequencing of the bacterial community with longer read technology would allow strain diversity identification and virome sequencing are required. In addition, denser longitudinal sampling is helpful to understand this dynamic with more details.

## 5 Data and Code Availability

The metagenomic raw data are available on GigaDB: http://gigadb.org/dataset/100145. The codes, needed files and SI are available on Github: https://github.com/JiqiuWu999/CRISPR/tree/master

## 6 Supplementary Information

The online version of this article contains supplementary material, which is available to authorized users.

## 7 Acknowledgements

With many thanks to Hanyun Zhang for coding help, Yan Xia for the data downloading and processing.

## 10 Supplementary Figures

**Supplementary Fig 1.**
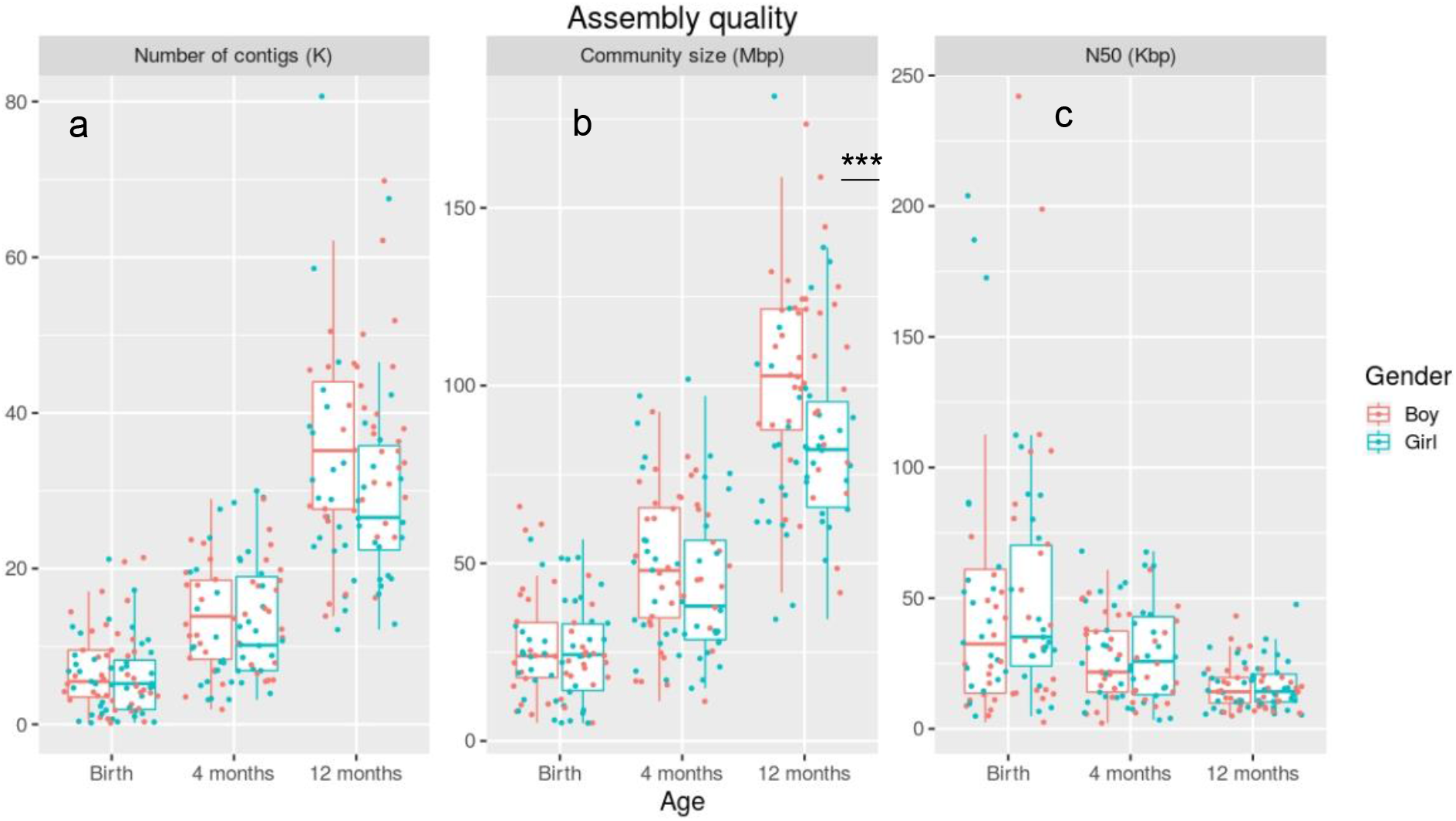
The quality of assemblies. (a) The number of contigs (K), (b) the community size (Mbp) and (c) the N50 (Kbp) in one sample. The difference in community size between girls and boys at 12 months was significant (Mann-Whitney U test, W = 1195, P = 0.00891).

**Supplementary Fig 2.**
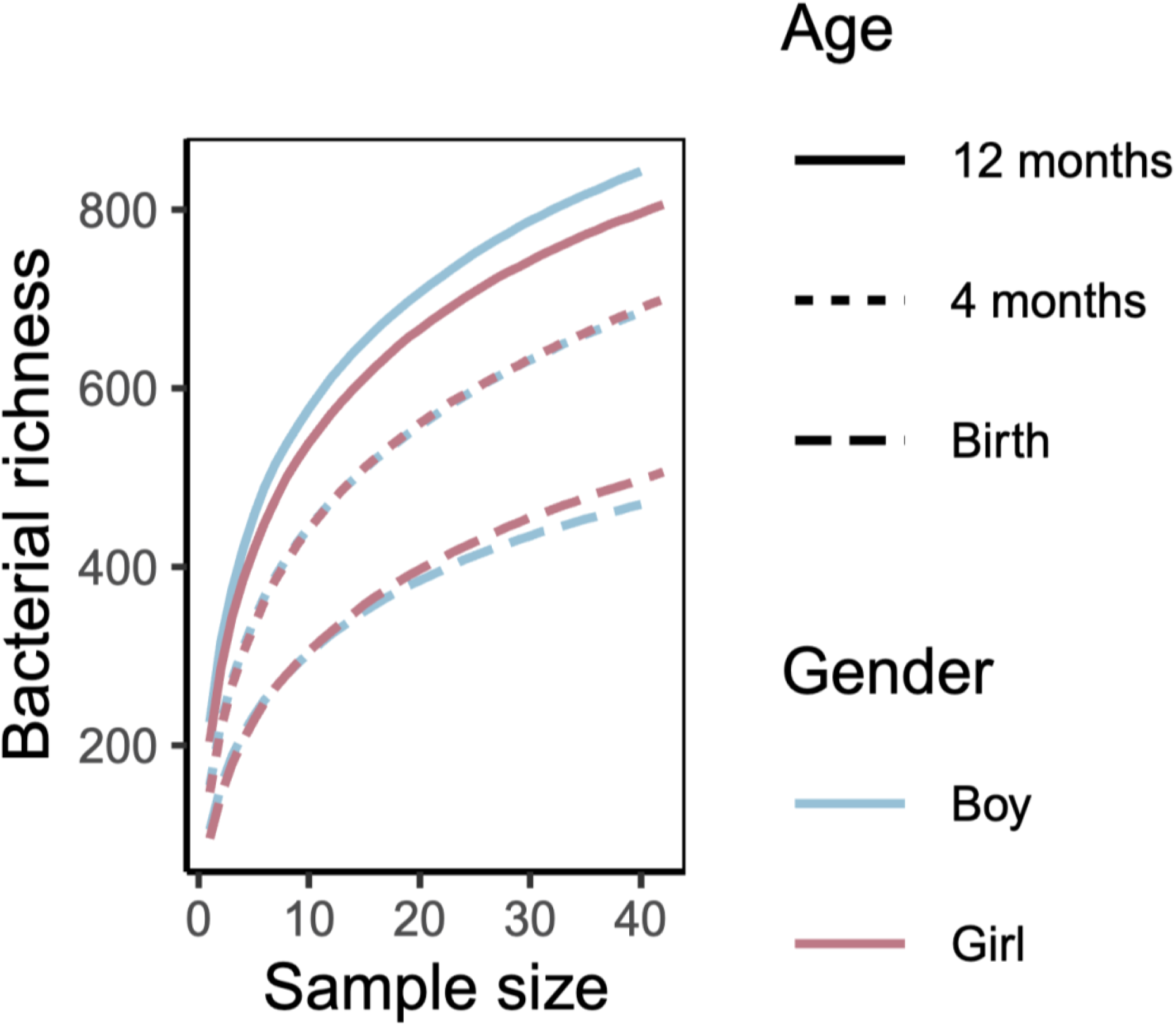
Rarefaction curve of bacterial richness.

**Supplementary Fig 3.**
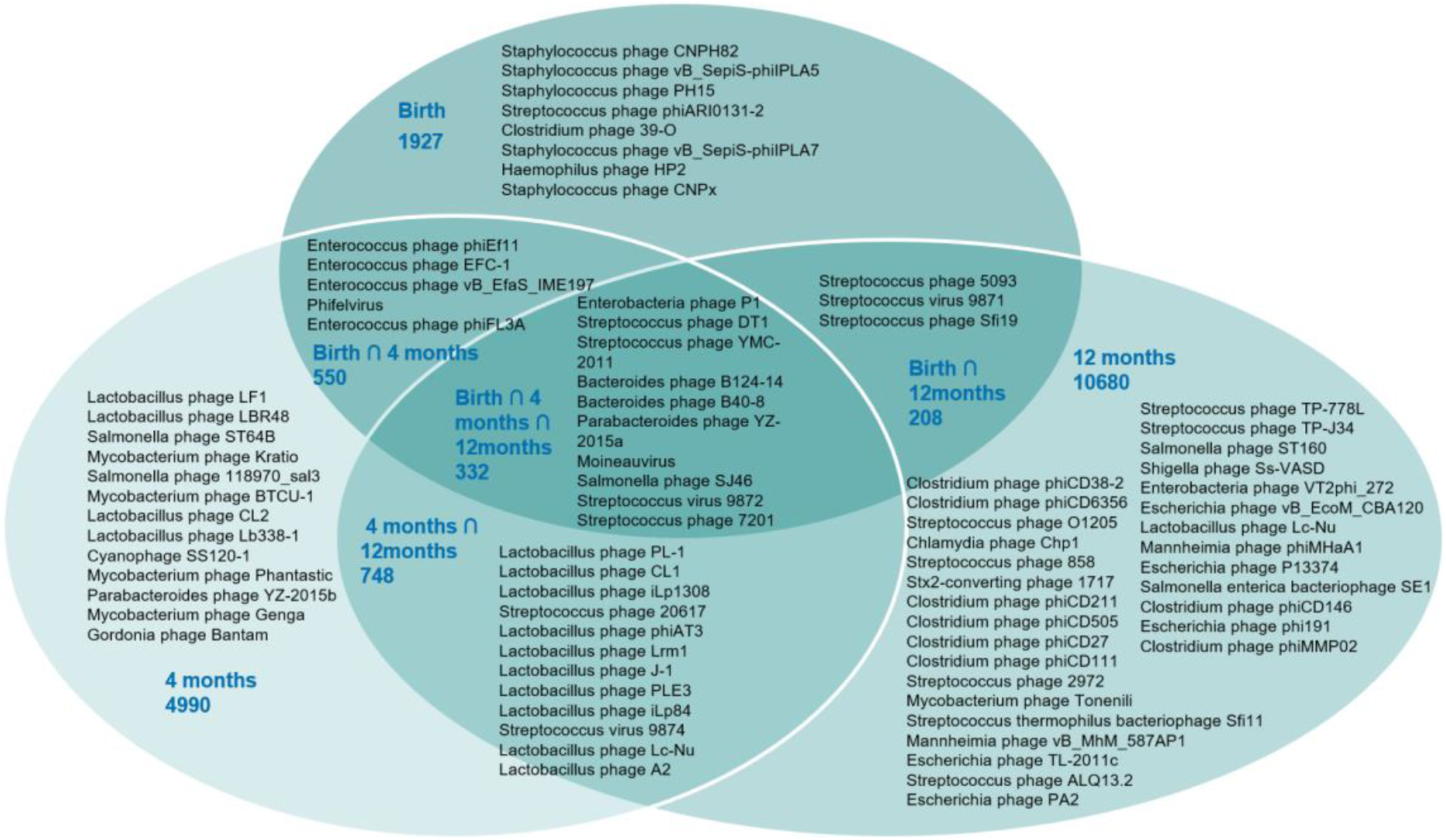
Bacteriophages targeted by the spacers. The number of unique and shared spacers at different time points and the bacteriophages targeted by the spacers. Only a small part of spacers was shared, and the bacteriophages targeted by spacers were alternated.

**Supplementary Fig 4.**
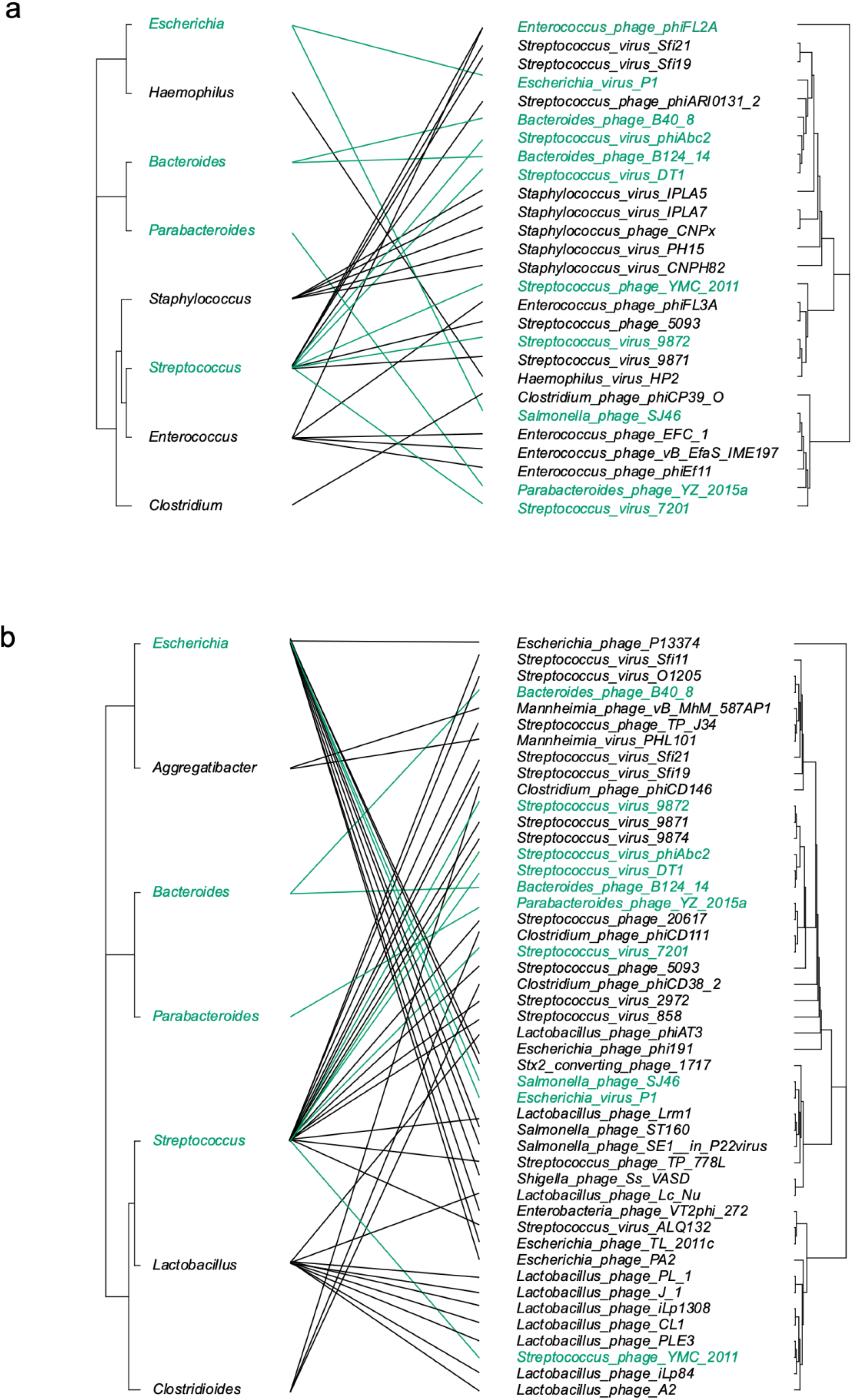
Spacers targeted distinct phages at birth and 12 months old. The interaction tree between bacteria and bacteriophages at birth (a) and 12 months old (b). The left and the right were the phylogenetic trees of bacteria and bacteriophages, respectively.

**Supplementary Fig 2.**
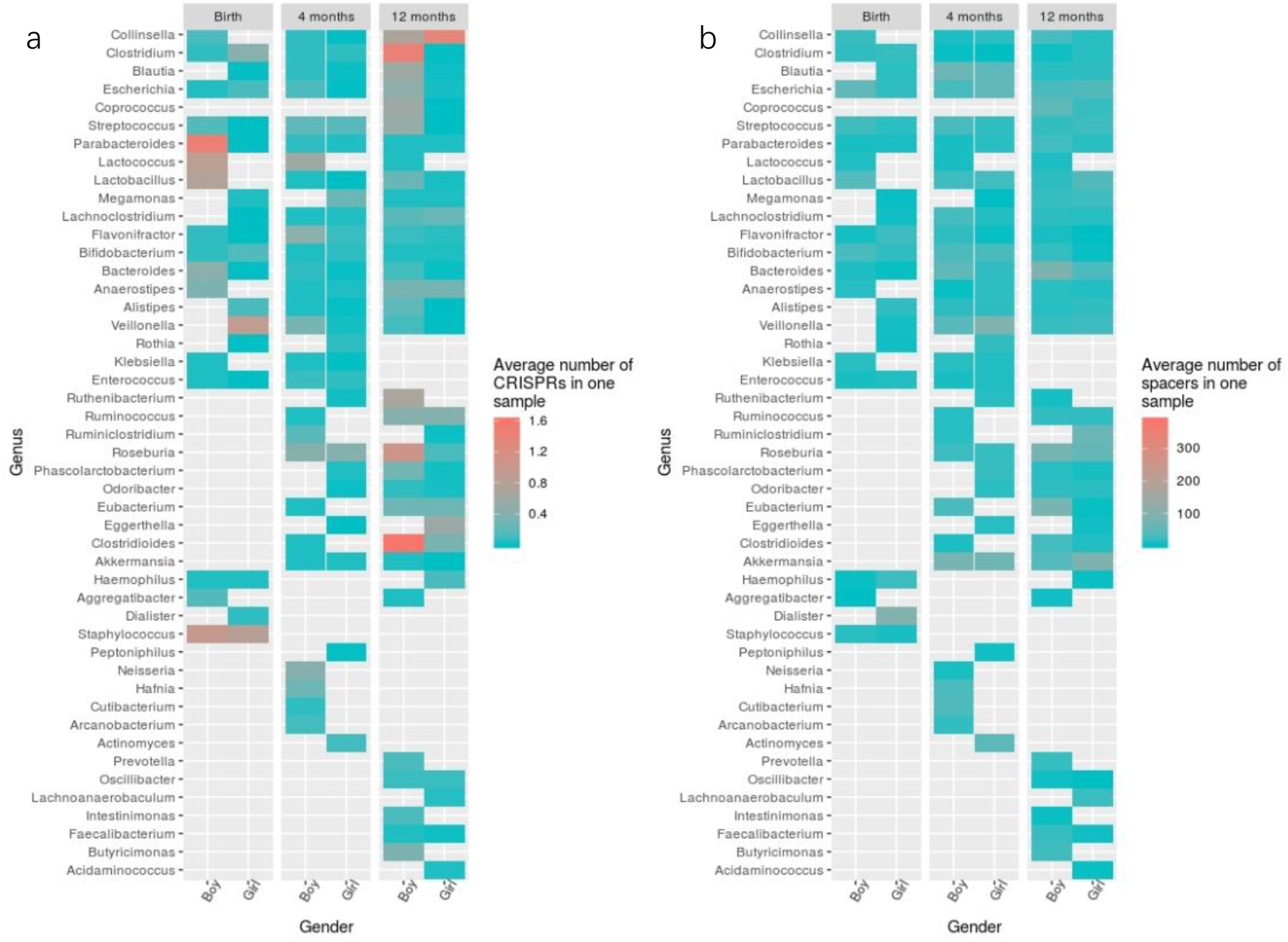
Boys contained more CRISPRs and spacers than girls in their gut communities. This heatmap showed the average number of CRISPRs (a) and spacers (b) in these representative bacteria in one sample. Overall, boys had more CRISPRs and spacers, however, the significant different of average number of CRISPRs and spacers in one bacterial species and in one sample between boys and girls was not observed.

